# ClC-7 links PIKfyve inhibition to Rab-dependent LRRK2 activity at lysosomes

**DOI:** 10.1101/2025.11.19.689251

**Authors:** Devin Clegg, Amanda Bentley-DeSousa, Agnes Roczniak-Ferguson, Shawn M. Ferguson

## Abstract

Increased activity of leucine-rich repeat kinase 2 (LRRK2) confers Parkinson’ s disease risk. LRRK2 dynamically localizes to lysosomal membranes in response to various stresses, yet the mechanisms by which distinct lysosomal perturbations are communicated to LRRK2 remain unclear. Here, we show that inhibition of the lysosomal lipid kinase PIKfyve promotes LRRK2 recruitment and signaling through a pathway that requires the lysosomal chloride/proton antiporter ClC-7. ClC-7 in turn controls the accumulation of multiple Rab GTPases on lysosomes. LRRK2 signaling under these conditions requires its established Rab-binding surfaces, with Rab12 contributing significantly to this response. This pathway operates independently of CASM. In contrast, lysosomal stresses that induce CASM require both Rab-binding sites on LRRK2 and GABARAP for robust LRRK2 signaling. These findings identify ClC-7-dependent lysosomal remodeling and Rab accumulation as key features linking PIKfyve inhibition to LRRK2 signaling and reveal that distinct lysosomal stresses engage different combinations of Rab and GABARAP inputs to activate LRRK2.

**Significance Statement:** This study defines how distinct lysosomal stresses activate LRRK2. PIKfyve inhibition engages a ClC-7-dependent, Rab-dependent pathway independently of GABARAP, whereas CASM-inducing stresses require both Rab interactions and GABARAP, revealing distinct mechanisms that couple lysosomal perturbations to LRRK2 signaling.

## Introduction

Mutations that increase signaling by leucine-rich repeat kinase 2 (LRRK2) are a major cause of familial Parkinson’s disease, and common LRRK2 variants also elevate risk for sporadic Parkinson’s disease (Alessi and Pfeffer, 2024; Bentley-DeSousa et al., 2025a; Paisan-Ruiz et al., 2004; Rocha et al., 2022; Zimprich et al., 2004). Pathogenic mutations in other Parkinson’s disease genes, including Rab32 and VPS35, also enhance LRRK2 signaling, underscoring the importance of understanding how upstream cellular processes regulate LRRK2 (Hop et al., 2024; McCarron et al., 2024; Mir et al., 2018). Defining these mechanisms is essential not only for explaining how inappropriate LRRK2 signaling contributes to Parkinson’s disease but also for understanding normal physiological functions of LRRK2.

Recent studies have revealed that LRRK2 signaling is strongly stimulated by lysosomal stress, leading to its recruitment to lysosomal membranes, increased kinase activity and phosphorylation of Rab GTPases (Bentley-DeSousa et al., 2025a; Bentley-DeSousa et al., 2025b; Bonet-Ponce et al., 2020; Kalogeropulou et al., 2020; McCarron et al., 2024). Lysosomal membrane damage and other triggers of proton efflux promote LRRK2 signaling through the process known as conjugation of ATG8 proteins to single membranes (CASM), which enables GABARAP proteins to recruit LRRK2 to stressed lysosomes (Bentley-DeSousa et al., 2025b; Eguchi et al., 2024; Huang et al., 2025).

In addition to GABARAP, multiple Rab GTPases also bind directly to distinct sites on LRRK2 and promote its recruitment to membranes and downstream signaling (Alessi and Pfeffer, 2024; Dhekne et al., 2023; Vides et al., 2022; Wang et al., 2023). Yet, how these Rab-dependent and GABARAP-dependent mechanisms of LRRK2 regulation are coordinated at lysosomes, and which specific forms of lysosomal stress favor one mode of regulation over the other, remain unresolved. This gap limits our understanding of how LRRK2 interprets the diverse stress signatures that arise both during normal lysosome homeostasis and under pathological conditions linked to Parkinson’s disease.

The lipid kinase PIKfyve synthesizes phosphatidylinositol-3,5-bisphosphate [PI(3,5)P₂], a low-abundance signaling lipid essential for endolysosomal membrane dynamics and ion homeostasis (Gary et al., 1998; Leray et al., 2022; Rivero-Rios and Weisman, 2022). Inhibition of PIKfyve depletes PI(3,5)P₂ and thus affects the activities of ion channels and other proteins that bind this lipid. A major target of PI(3,5)P_2_ is the chloride/proton (Cl⁻/H⁺) antiporter known as ClC-7. As PI(3,5)P_2_ inhibits ClC-7, inhibition of PIKfyve causes lysosomal chloride influx (Cao et al., 2023; Leray et al., 2022; Nicoli et al., 2019; Wu et al., 2025). This results in massive enlargement of lysosomes, with Cl^-^ acting as a counter-ion that supports further lysosome acidification, endosome-lysosome fusion as well as the potential delivery of new membrane via contact sites with the endoplasmic reticulum (Choy et al., 2018; Leray et al., 2022; Yang et al., 2025). Previous work linked PIKfyve inhibition to increased LRRK2 activity in the context of other lysosomal perturbations, including LLOME, nigericin, and chloroquine (McCarron et al., 2024). This observation was important because it placed PIKfyve inhibition within the expanding set of lysosome-associated triggers of LRRK2 signaling. However, it also highlighted an unresolved conceptual problem: these perturbations do not impose the same kind of lysosome stress. Unlike CASM-inducing stimuli that cause lysosome membrane damage and/or deacidification, PIKfyve inhibition depletes PI(3,5)P₂, activates ClC-7-dependent lysosomal chloride influx, promotes lysosome acidification, and drives lysosome enlargement. These observations argued against a universal CASM-dependent mechanism for LRRK2 activation. This problem further intersects with unanswered questions concerning Rab- and GABARAP-dependent LRRK2 regulation.

Here, we investigated LRRK2 signaling across mechanistically distinct lysosomal stresses to determine how Rab and GABARAP interactions are integrated at lysosomes. We show that PIKfyve inhibition activates LRRK2 through a CASM-independent pathway that requires ClC-7 and is accompanied by the accumulation of multiple Rab GTPases on enlarged lysosomes. Activation of LRRK2 signaling further requires its established Rab-binding surfaces, with Rab12 contributing to this response, but does not require CASM or GABARAP. Together, these findings argue that LRRK2 activation is not governed by a universal lysosome-stress pathway, but by the integration of distinct lysosomal state cues that engage combinations of Rab- and GABARAP-dependent mechanisms to control LRRK2 signaling.

## Results and Discussion

### PIKfyve inhibition activates LRRK2 and drives its recruitment to lysosomes

Pharmacological inhibition of PIKfyve with apilimod causes pronounced lysosome enlargement across multiple cell types, including the Raw264.7 mouse macrophage cell line (Choy et al., 2018; Gayle et al., 2017; Rivero-Rios and Weisman, 2022). We observed that this effect of apilimod on Raw264.7 cells was accompanied by a robust increase in Rab10 phosphorylation at threonine 73, a well-established direct LRRK2 substrate, which was similarly induced by additional PIKfyve inhibitors including vacuolin-1 and APY0201 (Fig. 1A-B, S1A-S1D) (Alessi and Pfeffer, 2024). Elevated phosphorylation of additional LRRK2 substrates, Rab8A (T72) and Rab12 (S106), further confirmed the increase in LRRK2 kinase activity (Fig. S1E–H). Note that as this phospho-Rab8 antibody cross-reacts with other Rabs that are LRRK2 substrates, this antibody may better reflect a pan-phospho-Rab readout rather than just the status of Rab8A-T72 (Lis et al., 2018). The absence of Rab10 phosphorylation in LRRK2-knockout (KO) cells and its loss upon treatment with the LRRK2 inhibitor MLi-2 further demonstrated LRRK2 dependence of Rab phosphorylation changes downstream of PIKfyve inhibition (Fig. 1A,1E-F). Immunoblot analyses of purified lysosomes revealed accumulation of LRRK2, Rab10 and phospho-Rab10 at lysosomes following PIKfyve inhibition (Fig. 1G–I). Comparable LRRK2 signaling in response to PIKfyve inhibition was observed in mouse embryonic fibroblasts, the mouse BV-2 microglia cell line, and human iPSC-derived microglia (Fig. S1I–L, 1J-K), indicating broad conservation across species and cell types. Furthermore, immunofluorescence analysis in both Raw264.7 cells and human iPSC-derived microglia demonstrated that a subset of enlarged lysosomes was the major site of phosphorylated Rab10 localization following apilimod treatment (Fig. 1L and M). These results are consistent with a close relationship between LRRK2 lysosome localization and Rab phosphorylation in macrophages and microglia (Bentley-DeSousa et al., 2025a).

**Figure 1.**
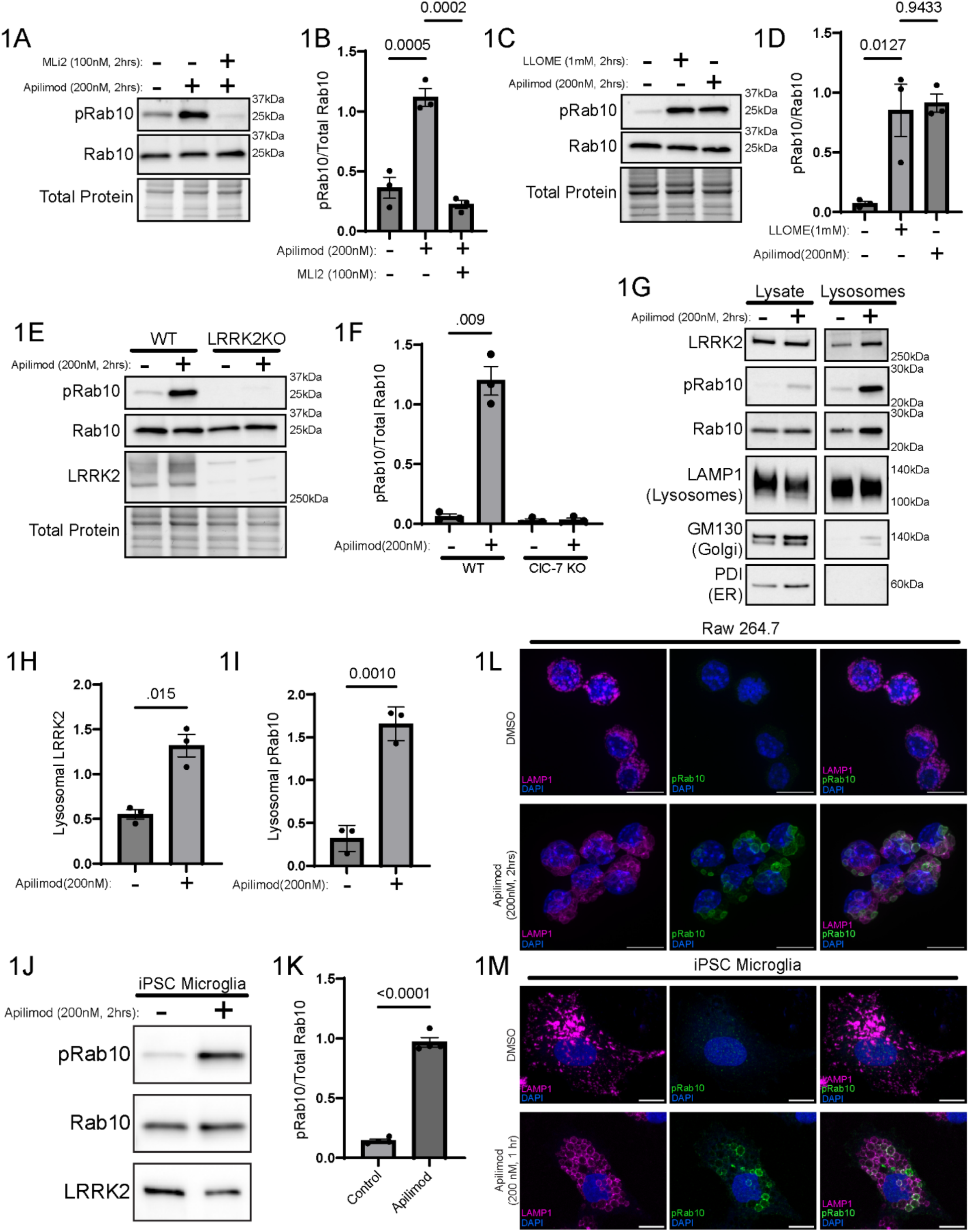
PIKfyve Inhibition Activates LRRK2 and Drives its Recruitment to Lysosomes. (A) Immunoblot of Raw264.7 cells treated with apilimod (200nM, 2 h) ± MLi-2 (100nM, 2 h) and probed for pRab10 (T73). (B) Quantification of pRab10 (T73) after apilimod ± MLi-2 (mean ± SEM; raw values, n = 3, one-way ANOVA, Sidak’s). (C) Immunoblot of cells treated with apilimod (200nM, 2 h) or LLOME (1mM, 2 h) and probed for pRab10 (T73). (D) Quantification of pRab10 (T73) after apilimod or LLOME treatment (mean ± SEM; raw values, n = 3, one-way ANOVA, Sidak’s). (E) Immunoblot of wild-type and LRRK2 KO cells treated with apilimod (200nM, 2 h) and probed for pRab10 (T73). (F) Quantification of pRab10 (T73) in wild-type and LRRK2 KO cells (mean ± SEM; raw values, n = 3, two-tailed unpaired Welch’s t test). (G) Immunoblot of lysates and purified lysosomes from cells treated with apilimod (200nM, 2 h). Images for the lysate and lysosome fractions come from the same exposure of the same blot that was loaded with 5µg lysate and 0.5µg of lysosome protein. (H) Quantification of lysosomal LRRK2 (mean ± SEM; raw values, n = 3, two-tailed unpaired Welch’s t test). (I) Quantification of lysosomal pRab10 (T73) (mean ± SEM; raw values, n = 3, two-tailed unpaired Welch’s t test). (J) Immunoblot of iPSC-derived microglia treated with DMSO or apilimod (200nM, 2 h) and probed for pRab10 (T73). (K) Quantification of pRab10 (T73) after apilimod treatment (mean ± SEM; raw values, n = 3, two-tailed unpaired Welch’s t test). (L) Immunofluorescence imaging of Raw264.7 cells treated with DMSO or apilimod (200nM, 2 h; LAMP1, magenta; pRab10 (T73), Green; DAPI, Blue). (M) Immunofluorescence imaging of iPSC-derived microglia treated with DMSO or apilimod (200nM, 2 h) (LAMP1, magenta; pRab10 (T73), Green; DAPI, Blue).

### ClC-7 is required for LRRK2 signaling downstream of PIKfyve inhibition

To investigate molecular mechanisms linking PIKfyve inhibition to LRRK2 signaling, we next focused on ClC-7, a major mediator of the cellular effects of PIKfyve inhibition whose activity is strongly enhanced when PI(3,5)P₂ levels fall (Gayle et al., 2017; Leray et al., 2022)(Fig. S2A). In ClC-7-KO cells, PIKfyve inhibition resulted in a more limited increase in lysosome size and did not significantly elevate Rab10 phosphorylation (Fig. 2A–D). LRRK2 accumulation at lysosomes was likewise impaired (Fig. 2E–F). TRPML1 is a lysosomal cation channel that is positively regulated by PI(3,5)P_2_ (Dong et al., 2010). However, pharmacological inhibition of TRPML1 had no significant effect on LRRK2 signaling (Fig. S2C-D). These observations collectively support a model in which ClC-7 is a key intermediate between PIKfyve and LRRK2.

**Figure 2.**
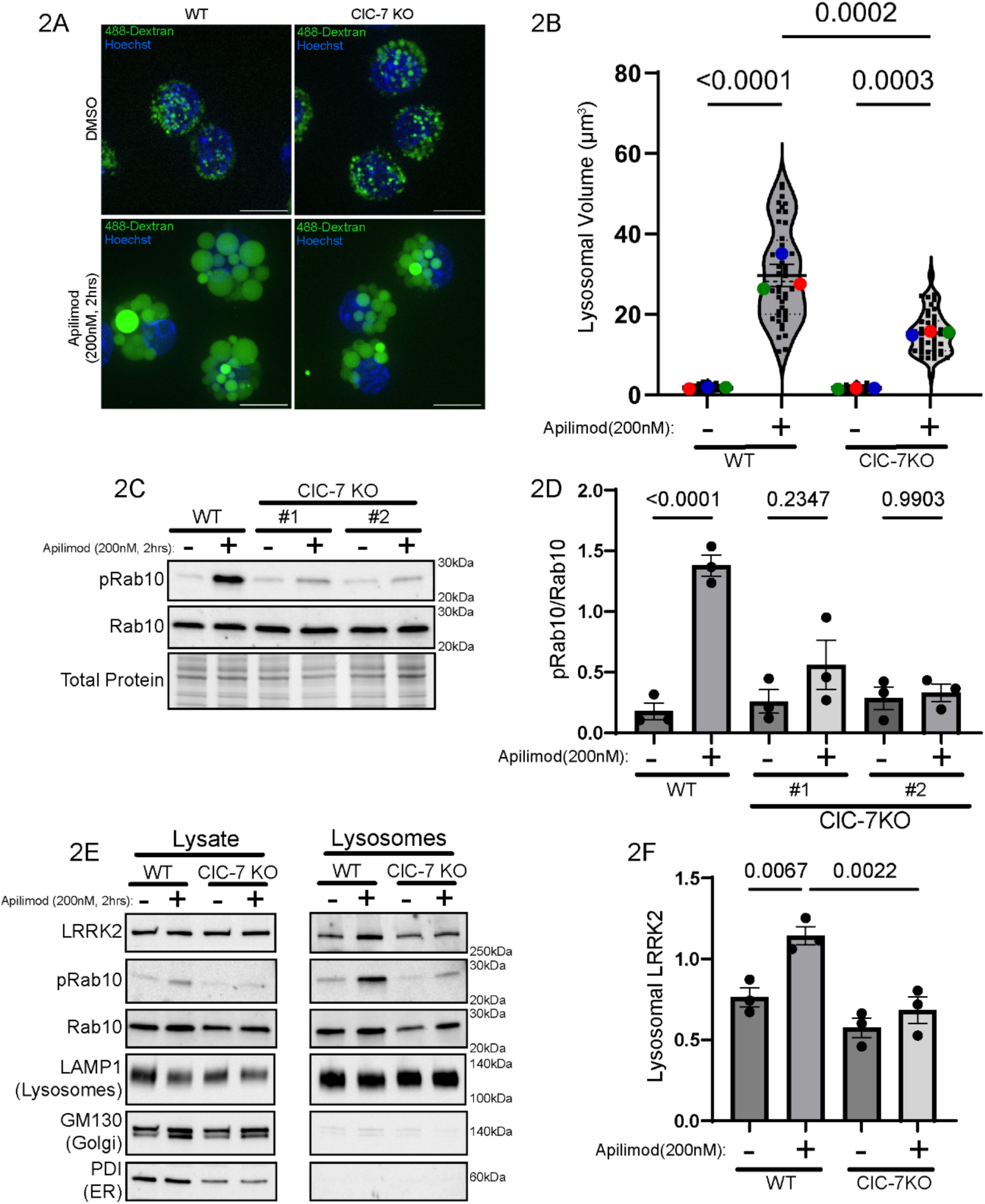
ClC-7 is required for LRRK2 activation by PIKfyve inhibition. (A) Live cell images of wild-type and ClC-7 KO Raw264.7 cells treated with DMSO or apilimod (200nM, 2 h) (488-Dextran, Green; Hoechst, Blue). (B) Quantification of lysosomal volume from wild-type and ClC-7 KO raw cells treated with DMSO and apilimod (200nM, 2 h) (mean ± SEM; raw values, n = 3, one-way ANOVA, Sidak’s). (C) Immunoblot of wild-type and ClC-7 KO cells treated with DMSO or apilimod and probed for pRab10 (T73). (D) Quantification of pRab10 (T73) from wild-type and ClC-7 KO cells treated with DMSO or apilimod (mean ± SEM; raw values, n = 3, one-way ANOVA, Sidak’s). (E) Immunoblot of lysates and purified lysosomes from wild-type and ClC-7 KO cells treated with apilimod (200nM, 2 h). Images for the lysate and lysosome fractions come from the same exposure of the same blot that was loaded with 10µg lysate and 1µg of lysosome protein. (F) Quantification of lysosomal LRRK2 (mean ± SEM; raw values, n = 3, one-way ANOVA, Sidak’s).

We next pharmacologically interfered with apilimod-induced lysosome enlargement using two complementary approaches. Inhibition of the V-ATPase with bafilomycin A1 prevented apilimod-induced lysosome enlargement and suppressed LRRK2 activation (Fig. S2E-G) (Leray et al., 2022). Likewise, treatment with phloretin, an inhibitor of water channels and/or other transporters required for osmotic swelling of lysosomes in response to apilimod, blocked both lysosome enlargement and the associated increase in LRRK2 kinase activity (Fig. S2E, H-I)(Yang et al., 2025).

### Rab interactions are required for LRRK2 recruitment and activation following apilimod treatment

We next sought to define protein-protein interactions that recruit LRRK2 to lysosomes and promote its signaling following PIKfyve inhibition. Although multiple Rab-binding sites on LRRK2 have been defined, upstream signals that promote Rab-dependent activation of LRRK2 remain incompletely understood (Alessi and Pfeffer, 2024; Bentley-DeSousa et al., 2025a; Dhekne et al., 2023; Wang et al., 2023). Because Rab GTPases can mediate LRRK2 membrane recruitment, we tested their requirement in apilimod-induced LRRK2 signaling. To this end, LRRK2-KO Raw264.7 cells were complemented with either wild-type LRRK2 or a Rab-binding-deficient mutant (K17/K18A + E240R + K439E, subsequently referred to “3×Rab mutant”) lacking the three major Rab-interaction sites that collectively support binding to Rab8A, 8B, 10, 12, 29, 32, and 38 (Alessi and Pfeffer, 2024; Bentley-DeSousa et al., 2025a)(Fig. 3A, S3A). The 3×Rab mutant failed to phosphorylate Rab substrates or accumulate on lysosomes following PIKfyve inhibition (Fig. 3B-E). These experiments collectively establish the importance of LRRK2-Rab interactions via sites 1, 2 and 3 as critical for LRRK2 signaling from lysosomes following PIKfyve inhibition.

**Figure 3.**
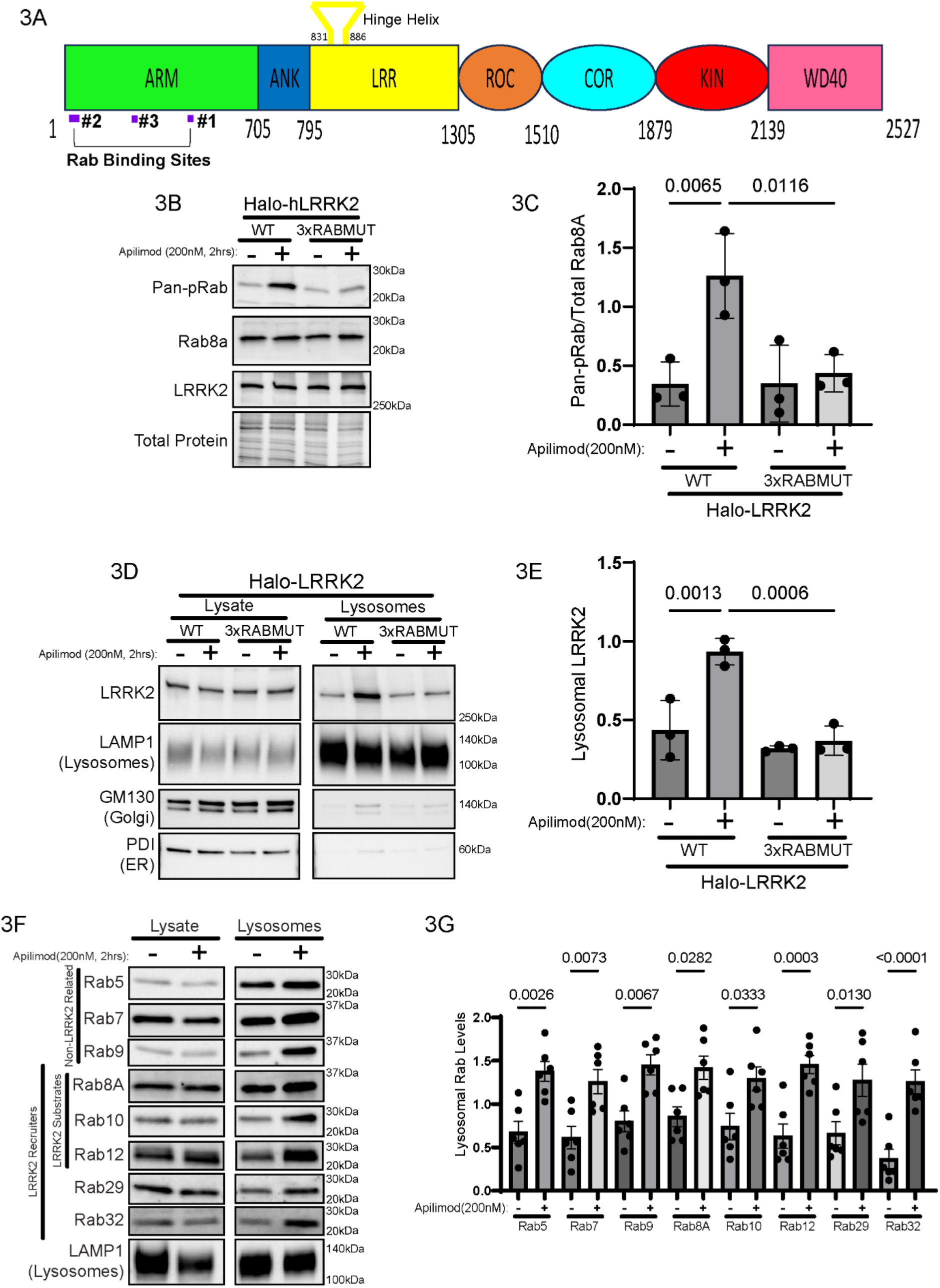
Rab GTPase interactions are required for LRRK2 recruitment and activation. (A) Sequence of LRRK2 showing its three Rab-binding sites. (B) Immunoblot of Raw264.7 cells stably expressing wild-type or Rab-binding-deficient (3xRABMUT) LRRK2 treated with apilimod (200nM, 2 h) and probed for Pan-pRab. (C) Quantification of Pan-pRab in wild-type and 3xRABMUT cells treated with apilimod (mean ± SEM; raw values, n = 3, one-way ANOVA, Sidak’s). (D) Immunoblot of lysates and lysosomes from Raw 264.7 cells stably expressing wild-type or Rab-binding-deficient (3xRABMUT) LRRK2 treated with apilimod (200nM, 2 h). Images for the lysate and lysosome fractions come from the same exposure of the same blot that was loaded with 10µg lysate and 1µg of lysosome protein. (E) Quantification of lysosomal LRRK2 from wild-type and 3XRABMUT cells treated with apilimod (mean ± SEM; raw values, n = 3, one-way ANOVA, Sidak’s). (F) Immunoblot analysis of Rab GTPases (Rab5, Rab7, Rab8A, Rab9, Rab10, Rab12, Rab29, and Rab32) from lysates and purified lysosomes from Raw264.7 cells treated with apilimod (200nM, 2 h). Images for the lysate and lysosome fractions come from the same exposure of the same blot that was loaded with 10µg lysate and 1µg of lysosome protein. (G) Quantification of lysosomal Rab GTPases (Rab5, Rab7, Rab8A, Rab9, Rab10, Rab12, Rab29, and Rab32) levels (mean ± SEM; raw values quantified from immunoblots in Fig. 1G and 5A, n = 3, one-way ANOVA, Sidak’s).

### Distinct stress pathways converge on Rab and GABARAP-dependent LRRK2 activation

Having established a key role for Rab-dependent LRRK2 signaling in response to the lysosome stress arising from PIKfyve inhibition and ClC-7 activation, we next asked whether the requirement for ClC-7 activity and LRRK2-Rab interactions extended to other lysosomal perturbations known to promote LRRK2 signaling, including those that trigger CASM and GABARAP lipidation. We therefore compared the ClC-7–dependent pathway with the CASM– GABARAP pathway, which functions in response to a variety of signals including lysosome membrane damage or deacidification (Bentley-DeSousa et al., 2025a; Bentley-DeSousa et al., 2025b). Stimuli that engage CASM—including LLOME and nigericin—continued to activate LRRK2 in ClC-7 KO cells (Figs. 4A-B), confirming that distinct upstream mechanisms can independently trigger the kinase. However, PIKfyve-induced activation was unaffected by perturbations that disrupt CASM-dependent LRRK2 activation, including ATG16L1 knockout, expression of the CASM inhibitor SopF, or GABARAP knockout (Fig. 4C-H). Conversely, even though CASM-GABARAP was not required for LRRK2 signaling that was triggered by PIKfyve inhibition, the Rab-binding–deficient LRRK2 mutant failed to signal in response to CASM-inducing stimuli (LLOME, nigericin, or DMXAA) (Figs. 4I-L).

**Figure 4.**
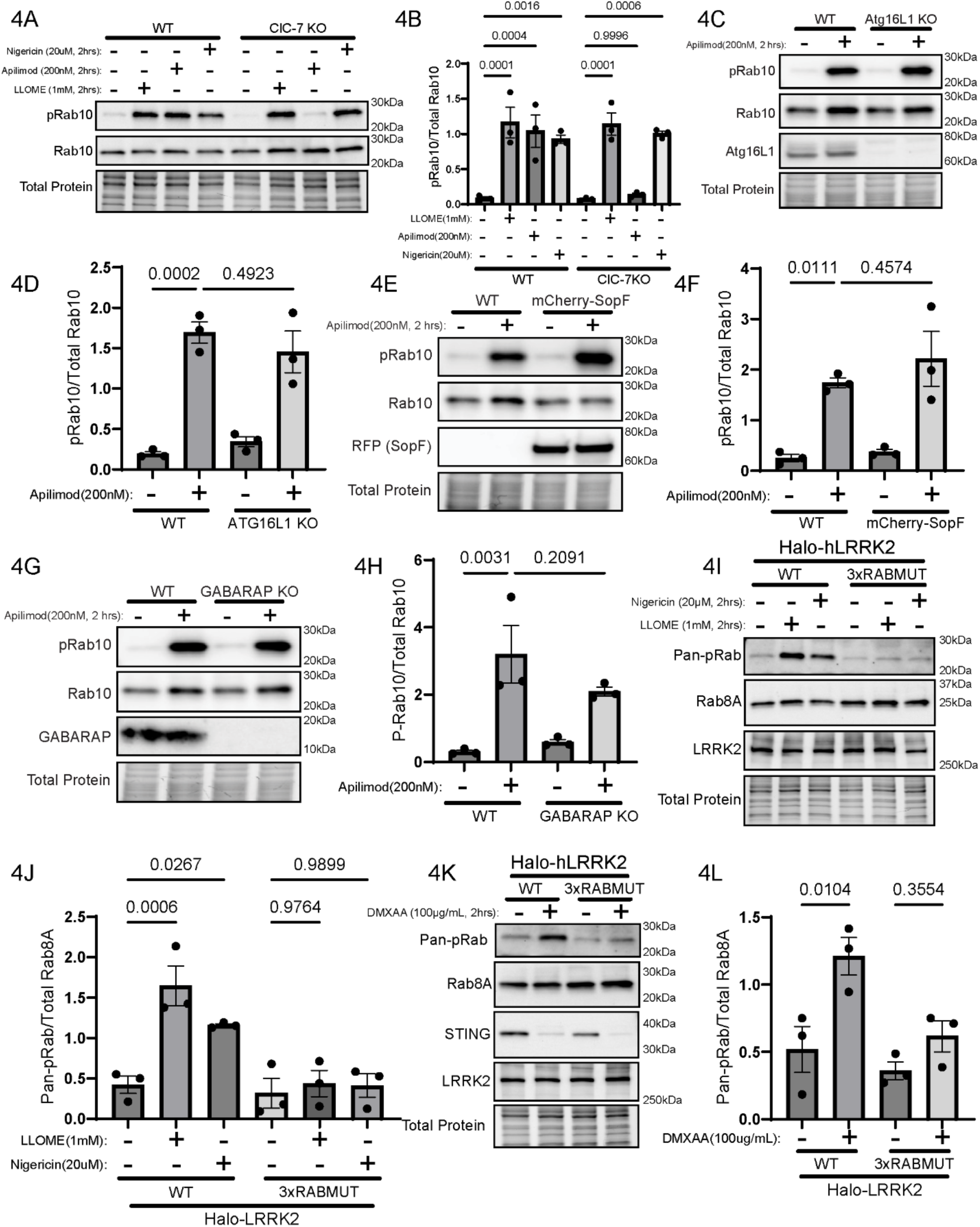
LRRK2 Activation Following Distinct Lysosome Stressors Converges on LRRK2-Rab GTPase Interactions. (A) Immunoblots of wild-type and ClC-7 KO cells treated with apilimod (200nM, 2 h) or CASM activators (LLOME, nigericin; 1mM, 20uM for 2 h). (B) Quantification of pRab10 (T73) after apilimod or CASM activators (mean ± SEM; raw values, n = 3, two-tailed unpaired Welch’s t test). (C) Immunoblots of wild-type and ATG16L1 KO cells treated with DMSO or apilimod (200nM, 2 h). (D) Quantification of pRab10 (T73) from apilimod-treated wild-type and ATG16L1 KO cells (mean ± SEM; raw values, n = 3, one-way ANOVA, Sidak’s). (E) Immunoblots of wild-type and mCherry-SopF-expressing cells treated with DMSO or apilimod (200nM, 2 h). (F) Quantification of pRab10 (T73) from apilimod-treated wild-type and mCherry-SopF-expressing cells (mean ± SEM; raw values, n = 3, one-way ANOVA, Sidak’s). (G) Immunoblots of wild-type and GABARAP KO cells treated with DMSO or apilimod (200nM, 2 h). (H) Quantification of pRab10 (T73) from apilimod-treated wild-type and GABARAP KO cells. (mean ± SEM; raw values, n = 3, one-way ANOVA, Sidak’s). (I) Immunoblot of Raw 264.7 cells stably expressing wild-type or Rab-binding-deficient (3xRABMUT) LRRK2 treated with LLOME (1mM, 2 h) or Nigericin (20µM, 2 h). (J) Quantification of Pan-pRab in wild-type and 3xRABMUT cells treated with LLOME or Nigericin (mean ± SEM; raw values, n = 3, one-way ANOVA, Sidak’s). (K) Immunoblot of Raw 264.7 cells stably expressing wild-type or Rab-binding-deficient (3xRABMUT) LRRK2 treated with DMXAA (100µg/mL, 2 h). (L) Quantification of Pan-pRab in wild-type and 3xRABMUT cells treated with DMXAA (mean ± SEM; raw values, n = 3, one-way ANOVA, Sidak’s).

The requirement for Rab interactions in promoting LRRK2 signaling raised questions about the identity of critical Rabs and their responses to lysosome stresses. However, this is a challenging problem to solve given the large number of Rabs that interact with LRRK2 via sometimes overlapping binding sites. To begin to narrow down the list of candidates, we first investigated which Rabs accumulate on lysosomes in response to PIKfyve inhibition. We were surprised to find that Rabs 5, 7, 8A, 9, 10, 12, 29 and 32 all increased in their abundance at lysosomes after apilimod treatment (Fig. 3F-G, 5A). Note that this included both Rabs that bind LRRK2 (8A, 10, 12, 29 and 32), Rabs that are LRRK2 substrates (8A, 10, 12) and even Rabs that are neither LRRK2 binding partners nor substrates (5, 7 and 9). To further narrow the list, we next reasoned that given the requirement for ClC-7 in apilimod-dependent stimulation of LRRK2 lysosome localization and signaling, ClC-7 KO cells should lose recruitment of Rabs that are critical for LRRK2 signaling. We found that Rabs 5, 9, 12 and 32 exhibited reduced apilimod-induced lysosome accumulation in the ClC-7 KO (Fig. 5A-B). Given that Rabs 12 and 32 are well established LRRK2-interacting proteins, these results suggested important roles for Rab12 and Rab32 in LRRK2 regulation by this stimulus (Dhekne et al., 2023; McGrath et al., 2021; Wang et al., 2023). We tested this by using CRISPR to deplete Rab12 in combination with siRNA-mediated depletion of Rab32 (Fig. 5C). Loss of Rab12 significantly reduced the phosphorylation of Rab10 induced by apilimod treatment while the Rab32 knockdown had no effect either on its own or in combination with Rab12 depletion. Collectively, these results point to an important role for Rab12 in supporting LRRK2 signaling downstream of PIKfyve inhibition. This is in line with a similar impact of Rab12 KO on apilimod-induced LRRK2 signaling that was recently reported elsewhere (Malankhanova et al., 2025).

**Figure 5.**
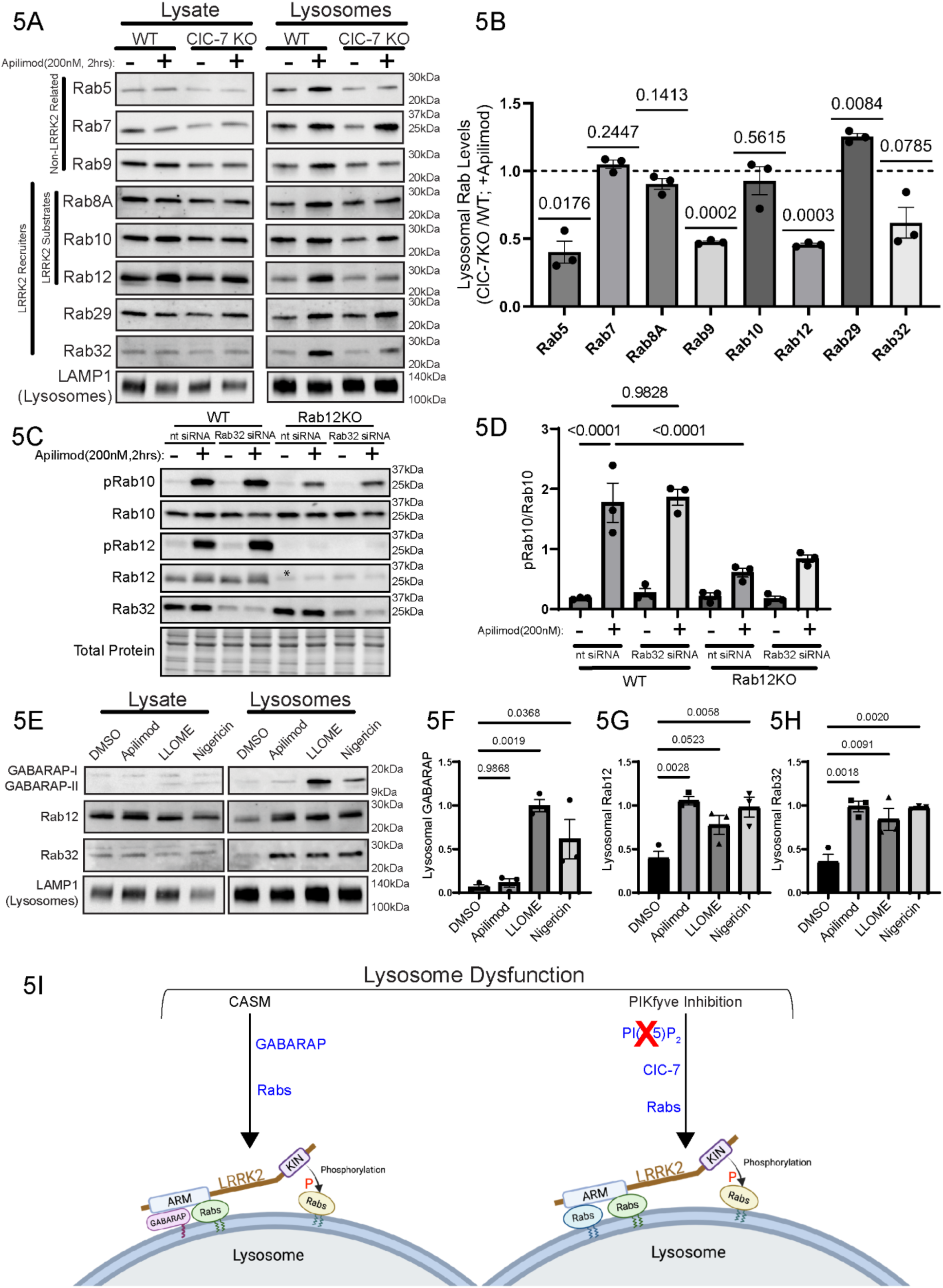
Distinct Lysosome Stresses Converge on LRRK2 Membrane Recruitment and Activation through Accumulation of Lysosomal GABARAP and Rab GTPases. (A) Immunoblots of lysates and lysosomes from wild-type and ClC-7 KO cells +/- the indicated treatments with apilimod (200nM, 2 hrs). Images for the lysate and lysosome fractions come from the same exposure of the same blot that was loaded with 10µg lysate and 1µg of lysosome protein. (B) Ratio of Lysosomal Rab GTPase levels in apilimod treated ClC-7 KO versus wild-type cells (ClC-7 KO/WT). (mean ± SEM; raw values, n = 3, one sample t test, hypothetical value = 1). (C) Immunoblots of WT and Rab12 KO Raw264.7 cells transfected with either non-targeting or Rab32 siRNA and then treated with DMSO or Apilimod (200nM, 2 h). (* denotes a non-specific band recognized by the Rab12 antibody) (D) Quantification of pRab10 levels in response to apilimod treatment +/- Rab12 and Rab32 genetic perturbations. (E) Immunoblots of lysates and lysosomes from apilimod (200nM, 2 h), LLOME (1mM, 2 h), or Nigericin (20µM, 2 h) treated wild-type cells. Images for the lysate and lysosome fractions come from the same exposure of the same blot that was loaded with 10µg lysate and 1µg of lysosome protein. The lysate GABARAP blot is shown at a longer exposure. (F) Quantification of lysosomal GABARAP levels in wild-type cells treated with apilimod, LLOME, or Nigericin (mean ± SEM; raw values, n = 3, one-way ANOVA, Sidak’s). (G) Quantification of lysosomal Rab12 levels in wild-type cells treated with apilimod, LLOME, or Nigericin (mean ± SEM; wild-type, n = 3, one-way ANOVA, Sidak’s). (H) Quantification of lysosomal Rab32 levels in wild-type cells treated with apilimod, LLOME, or Nigericin (mean ± SEM; raw values, n = 3, one-way ANOVA, Sidak’s). (I) Model of how distinct lysosomal stresses use combinations of Rabs or GABARAP to converge on LRRK2 recruitment and signaling at lysosomes. Created with BioRender (Clegg, D. and Ferguson, S.M., 2026).

To further examine the relationship among lysosome stresses, CASM-GABARAP and Rabs upstream of LRRK2, we compared cell lysates and purified lysosomes from cells treated in parallel with apilimod, LLOME and nigericin. Consistent with the lack of requirement for GABARAP lipidation in the stimulation of LRRK2 signaling by apilimod (Fig. 5E-F), apilimod failed to trigger GABARAP lipidation at lysosomes while both LLOME and nigericin triggered robust responses. Meanwhile, all three stimuli resulted in an increase in the lysosomal abundance of Rabs 12 and 32 (Fig. 5E, G-H).

The observed overlap between LRRK2 “Substrate Rabs” (including Rabs 8, 10, 12) and LRRK2 “Recruitment Rabs” (Rabs 8A/B, 10, 12, 29, 32 and 38) raised the possibility that at least some of the Rab 8, 10 and 12 accumulation that we observed at lysosomes of apilimod-treated cells resulted from their LRRK2-mediated phosphorylation. This prediction was based on previous studies proposing that LRRK2-mediated phosphorylation could increase the abundance of membrane-associated Rabs by decreasing their extraction by Rab GDI (Eguchi et al., 2018; Pfeffer, 2022; Steger et al., 2016). We tested this by treating cells with apilimod alone or with apilimod plus MLi-2 to inhibit the kinase activity of LRRK2 and then analyzing the abundance of each of these proteins in cell lysates and purified lysosomes (Fig. S3C-G). We found that while MLi-2 effectively suppressed LRRK2-mediated Rab10 phosphorylation, it did not significantly reduce the lysosomal abundance of LRRK2, Rab 8A, Rab 10 or Rab12 at baseline or following apilimod treatment. These results argue against either a need for LRRK2-dependent phosphorylation to promote lysosomal Rab accumulation or the existence of a positive feedback loop between phosphorylated Rabs and LRRK2 recruitment to lysosomes (via Site 2) under these conditions.

### Rab and GABARAP interactions allow LRRK2 to respond to distinct lysosomal stresses

Taken together, these findings provide a basis for considering how PIKfyve inhibition, ClC-7 activity, and changes in lysosomal Rab composition are mechanistically linked to LRRK2 activation. When PI(3,5)P₂ levels fall following PIKfyve inhibition, disinhibition of ClC-7 drives lysosomal chloride influx that is accompanied by changes in acidification, fission, fusion, and lipid delivery from the endoplasmic reticulum via bridge-like lipid transport proteins (Choy et al., 2018; Leray et al., 2022; Yang et al., 2025). We observed that this response was accompanied by the accumulation of multiple Rab GTPases on lysosomes, including the known LRRK2 binding partners Rab8A, Rab10, Rab12, Rab29, and Rab32. Our results support a model in which these changes promote LRRK2 recruitment and activation through its established Rab-binding sites without requiring CASM or GABARAP. Rab12 contributes significantly to LRRK2 signaling downstream of PIKfyve inhibition but does not fully account for the response, suggesting that additional Rabs may cooperate in LRRK2 activation. We speculate that the broad accumulation of Rabs creates a high-density Rab platform that favors LRRK2 recruitment and activation. The mechanism responsible for this Rab accumulation remains to be defined, but it does not appear to require LRRK2 kinase activity: apilimod increased the lysosomal abundance of Rabs that are not known LRRK2 substrates, including Rab5, Rab7, Rab9, and Rab32, while LRRK2 substrates such as Rab8A, Rab10, and Rab12 still accumulated on lysosomes when LRRK2 kinase activity was inhibited. In contrast to this GABARAP-independent response to PIKfyve inhibition, lysosomal stresses that engage CASM and LRRK2–GABARAP interactions also require LRRK2 Rab-binding sites for robust signaling. Thus, Rab-dependent mechanisms provide a broadly required input for LRRK2 activation at stressed lysosomes, whereas GABARAP acts as an additional, stimulus-selective scaffold under CASM-inducing conditions.

The relevant Rab- and GABARAP-binding surfaces lie within the N-terminal Armadillo, Ankyrin, and leucine-rich repeat regions of LRRK2, raising the possibility that these interactions do more than recruit LRRK2 to membranes (Alessi and Pfeffer, 2024; Bentley-DeSousa et al., 2025a). Recent structural work supports a model in which LRRK2 autoinhibition depends on docking of the N-terminal repeats against the catalytic core, whereas activation requires undocking of this region (Villagran Suarez et al., 2026). We therefore speculate that multivalent engagement of Rab GTPases and GABARAP on lysosomal membranes may help stabilize activation-compatible N-terminal conformations of LRRK2. In this way, lysosomal Rabs and GABARAP could act not only as recruitment factors, but also as membrane-associated ligands that favor conformational states permissive for Rab phosphorylation.

Several questions remain. First, it will be important to further define the contributions of additional Rabs (beyond Rab12), and the kinetics by which Rab accumulation is coupled to LRRK2 recruitment and activation. Second, the accumulation of multiple Rabs on stressed lysosomes suggests a broader remodeling of the lysosomal surface during stress whose mechanism remains to be deciphered. Because the Rabs that accumulate after lysosome stress do not share a common set of known GEFs or GAPs, their accumulation may reflect another regulatory mechanism, such as inefficient GDI-mediated extraction from stressed lysosomal membranes (Homma et al., 2021). Furthermore, these Rabs may recruit additional effectors involved in lysosome repair, remodeling, or damage signaling and thus play an important role in cellular responses to lysosome damage that extend beyond effects on LRRK2.

### Potential Relevance to Parkinson’s disease

Transient activation of LRRK2 at stressed lysosomes may support adaptive responses such as membrane remodeling, repair, or restoration of lysosome function (Bonet-Ponce et al., 2020; Herbst et al., 2020). Although we used acute chemical perturbations of lysosomes as experimental tools, the lysosomal responses that they produce are more broadly informative about how lysosomes may adapt in both physiological and pathophysiological contexts. For example, acute PIKfyve inhibition leads to disruption of core lysosome properties, including membrane delivery, fusion, fission, osmotic balance, and ion homeostasis. These features are continuously challenged as lysosomes receive cargo, remodel their membranes, and respond to nutrient or damage signals.

The dual requirement for LRRK2-Rab interactions and LRRK2-GABARAP interactions under CASM-inducing conditions may help restrict LRRK2 signaling to lysosomes experiencing specific stress-associated cues while limiting inappropriate activation at other Rab- or GABARAP-positive compartments. However, genetic or environmental conditions that elevate lysosomal Rab accumulation or chronically perturb lysosome ionic balance, osmotic state, or membrane integrity could shift this system toward persistent LRRK2 activation. Parkinson’s disease-associated mutations in Rab32 and VPS35 activate LRRK2, whereas VPS13C loss-of-function mutations impair repair of damaged lysosomes (Gustavsson et al., 2024; Hop et al., 2024; Mir et al., 2018; Wang et al., 2025).

The coupling of ClC-7 activity to LRRK2 further suggests that lysosomal ionic and osmotic homeostasis are important inputs into LRRK2 regulation. This connection aligns with evidence that other Parkinson’s disease-associated lysosomal ion channels and transporters, including TMEM175 and ATP13A2, influence lysosomal pH and ion flux (Cang et al., 2015; Riederer et al., 2026; van Veen et al., 2020). Environmental toxicants linked to Parkinson’s disease, such as trichloroethylene, may similarly converge on lysosome dysfunction and LRRK2 hyperactivation, further broadening the relevance of these stress-integration mechanisms beyond inherited genetic risk (De Miranda et al., 2021). Together, these observations suggest that LRRK2 may be engaged by lysosomal stress signatures arising from multiple Parkinson’s disease-relevant perturbations.

## Summary

Together, our results support a model in which distinct lysosomal stresses engage LRRK2 through overlapping but mechanistically distinct pathways. PIKfyve inhibition activates ClC-7 and promotes the accumulation of Rab GTPases at lysosomes, with Rab12 acting as a key contributor to LRRK2 recruitment and activation independently of GABARAP. In contrast, CASM-inducing stresses engage both Rab GTPases and GABARAP to promote LRRK2 signaling. These findings advance understanding of how changes in lysosomal lipid and ion homeostasis are integrated to control LRRK2 signaling from stressed lysosomes (Fig. 5I).

## Methods

### Raw264.7 cell culture

Raw264.7 cells were cultured in DMEM (11965-092; Thermo Fisher Scientific) supplemented with 10% FBS (16140-071; Thermo Fisher Scientific) and 1% penicillin/streptomycin (15140122; Thermo Fisher Scientific) at 37°C with 5% CO₂, following the supplier’s instructions (ATCC). Cells were passaged using CellStripper (356230; Corning).

### Stable cell line generation

Stable cell lines were generated using a PiggyBac transposase system. Briefly, 2.5 × 10⁵ cells were plated per well in a 6-well dish. The following day, cells were transfected with Lipofectamine 2000 (11668019; Invitrogen) according to the manufacturer’s protocol, using a 1:2 ratio of PiggyBac transposon plasmid to gene-of-interest plasmid (total 1 µg DNA) (Pantazis et al., 2022). After 48 h, the medium was replaced, and after 72 h, puromycin (A11138-03; Gibco) was added at 3.5 µg/mL for 48–72 h. Cells were then washed and returned to fresh medium for recovery. Clonal populations were obtained by plating single cells into 96-well dishes and expanded for screening by immunoblotting. A detailed protocol is available at https://dx.doi.org/10.17504/protocols.io.yxmvm9nr5l3p/v1. A custom human Halo-LRRK2 (Rab-binding-deficient) PiggyBac vector was obtained from VectorBuilder, and will be made available through Addgene: pPB-EF1A-HALO-hLRRK2; Addgene_229738, pPB-EF1A-HALO-hLRRK2 (Rab binding mutant/3xRabmut K439E, K17/18A, E240R);

### Genome-edited cell lines

*Clcn7 and Rab12* knockout (KO) Raw264.7 cells were generated using the Synthego CRISPR Gene Knockout V2 Mouse Kit. Briefly, 2.5 × 10⁵ cells were plated in a 6-well dish and transfected the next day with ribonucleoprotein complexes composed of recombinant Cas9 (Synthego CRISPR Gene Knockout V2) and gene-specific sgRNAs using Lipofectamine CRISPRiMAX (CMAX00003; Thermo Fisher Scientific). After 48 h, the medium was changed, and after 72 h, single cells were plated into 96-well dishes to establish clonal populations. Following clonal expansion, Rab12 KO clones were screened by immunoblot using Rab12 antibody. For CLCN7 KO clones, genomic DNA was extracted using QuickExtract, and PCR was performed using primers flanking the guide RNA cut sites. PCR products were resolved on 1% agarose gels to identify clones with large deletions and then were sequenced to further confirm gene disruption. Clone 1 contained 129 bp and 134 bp deletions; these deletions extended from within exon 5 into the downstream intron and removed 62 bp of coding sequence. Clone 2 contained a 133 bp deletion that removed all 71 bp of exon 5 together with adjacent flanking intronic sequence. A detailed protocol is available at https://dx.doi.org/10.17504/protocols.io.5jyl88wj6l2w/v1.

### iPSC-Microglia Differentiation

An established iPSC line (WTC11) that was engineered for efficient doxycycline-inducible microglia differentiation was kindly provided by Li Gan (Weill-Cornell) and differentiated via their established protocol (Drager et al., 2022).

### Generation of SPIONs

Superparamagnetic iron oxide nanoparticles (SPIONs) were prepared following an established protocol (https://dx.doi.org/10.17504/protocols.io.eq2lyn69pvx9/v1). Briefly, 10mL of 1.2 M FeCl₂ (220299; Sigma-Aldrich) and 10mL of 1.8 M FeCl₃ (157740; Sigma-Aldrich) were combined under stirring, followed by the slow addition of 10 ml 30% NH₄OH (320145; Sigma-Aldrich) over 5 min. The resulting particles were washed three times with 100 ml water, resuspended in 80mL 0.3 M HCl (9535; J.T. Baker), and stirred for 30 min. Dextran (4 g; D1662; Sigma-Aldrich) was then added and stirred for 30 min. The particles were dialyzed against H₂O for ≥2 days with multiple water changes, centrifuged at 26,900 g for 30 min to remove aggregates, and stored at 4°C.

### SPION-mediated lysosome purification

A 90% confluent 15-cm dish of Raw264.7 cells was split into 8, 10cm dishes. The following day, cells were incubated for 1 h with 10 ml of fresh medium containing 5% SPIONs and 5 mM HEPES (pH 7.4, 15630-080; Thermo Fisher Scientific). Dishes were washed twice with PBS and incubated in fresh medium for 2 h to allow SPION washout. All subsequent steps were performed on ice. Cells were washed, scraped in PBS, and pelleted at 300 g for 5 min at 4°C. Pellets were resuspended in 1mL ice-cold HB buffer (5 mM Tris base [AB02000-05000; American Bio], 250 mM sucrose [S0389; Sigma-Aldrich], 1 mM EGTA [E4378; Sigma-Aldrich], pH 7.4) supplemented with protease (cOmplete Mini, EDTA-free; Roche) and phosphatase inhibitors (PhosSTOP; Roche). Cells were homogenized with 50 strokes in a Dounce homogenizer (DWK Life Sciences Wheaton, 06-434) and centrifuged at 800 g for 20 min at 4°C. LS magnetic columns (130042401; Miltenyi Biotec) were equilibrated with 2.5 ml HB buffer on a QuadroMACS separator (130-091-051; Miltenyi Biotec). The supernatant was applied to the columns, and the flow-through was reapplied once. Columns were washed with 3mL HB buffer, removed from the magnet, and lysosomes were eluted in 2.5mL HB buffer into ultracentrifuge tubes. Eluates were centrifuged at 55,000 rpm for 10 min at 4°C using a Beckman-Coulter TLA-100.3 rotor. Pelleted lysosomes were resuspended in ∼50 µl HB buffer for immunoblotting. A detailed protocol is available at https://dx.doi.org/10.17504/protocols.io.e6nvwbk32vmk/v1.

### Immunoblotting

For immunoblot analysis, 1 × 10⁶ cells were seeded per well in a 6-well dish. The next day, cells were washed twice with ice-cold PBS (1.1 mM KH₂PO₄, 155.2 mM NaCl, 3 mM Na₂HPO₄) and scraped into 50 µl ice-cold lysis buffer (50 mM Tris base, 150 mM NaCl, 1% Triton X-100, 1 mM EDTA) supplemented with protease (cOmplete Mini, EDTA-free; Roche) and phosphatase inhibitors (PhosSTOP; Roche). Lysates were centrifuged at 14,000 rpm for 8 min at 4°C to remove insoluble material, and protein concentrations were determined using the Coomassie Plus Protein Assay (23236; Thermo Fisher Scientific). Lysate supernatants were mixed 1:1 with Laemmli buffer (80 mM Tris-HCl, pH 6.8, 25.3% glycerol, 2.67% SDS, bromophenol blue, and 6.187% freshly added β-mercaptoethanol) and heated at 95°C for 3 min. Typically, 20 µg protein per sample was resolved on 4–15% Mini-PROTEAN TGX Stain-Free precast gels (Bio-Rad) in standard Tris-glycine-SDS buffer. For lysosome fractions, 10 µg total lysate and 1 µg purified lysosome sample were loaded. Proteins were transferred to 0.45-µm nitrocellulose membranes (1620115; Thermo Fisher Scientific) at 100V for 60 min. Membranes were visualized using Bio-Rad stain-free imaging or Ponceau S, blocked in 5% nonfat milk in TBST (10 mM Tris base, 150 mM NaCl, 0.1% Tween-20), and incubated overnight at 4°C with primary antibodies diluted in 5% BSA/TBST. After three 5-min washes in TBST, membranes were incubated for 1 h at room temperature with HRP-conjugated secondary antibodies in 5% milk/TBST, washed again, and developed using either SuperSignal West Pico PLUS or SuperSignal West Femto substrates (Thermo Fisher Scientific). Images were acquired with a Bio-Rad ChemiDoc MP imaging system. Antibodies used are listed in the Key Resources Table. A detailed protocol is available at https://dx.doi.org/10.17504/protocols.io.5qpvo9bmdv4o/v1. Source data for all immunoblots will be deposited in the corresponding data repository.

### Immunofluorescence and imaging

For immunofluorescence, 2 × 10⁵ cells were plated on 12-mm glass coverslips (633029; Carolina Biological Supplies) in 6 well dishes. After treatment, cells were fixed in 4% paraformaldehyde (19202; Electron Microscopy Sciences) in sodium phosphate buffer (153.56 mM Na₂HPO₄, 53.63 mM NaH₂PO₄, pH 7.3) for 30 min at room temperature. Cells were washed three times for 5 min with PBS and permeabilized/blocked in 3% BSA and 0.1% saponin (S4521; Sigma-Aldrich) in PBS for 15 min. Primary antibodies were applied overnight at 4°C, followed by three PBS washes and incubation with fluorescent secondary antibodies for 1 h at room temperature in the dark. Cells were washed again, mounted on slides (12-550-143; Thermo Fisher Scientific) using ProLong Gold Antifade Mountant (P36935; Thermo Fisher Scientific), and stored at 4°C. Images were acquired with a Nikon Ti2-E inverted microscope equipped with a Yokogawa CSU-W1 SoRa spinning-disk super-resolution module using a 60× SR Plan Apo IR oil-immersion objective. Images were processed and adjusted in FIJI (version 2.14.0/1.54f). A detailed protocol is available at https://dx.doi.org/10.17504/protocols.io.36wgqd7wkvk5/v1.

### Live Cell Imaging

3.0 x 10^5^ cells were plated on Mattek dishes one day prior to the experiment. The next day, cells were incubated in media with Alexa Fluor 488 dextran (D22910;Invitrogen, 10mg/mL) 1:1000 dilution for 8 hours. Media was replaced to washout 488-dextran that was not endocytosed.and cells were treated with DMSO or apilimod (200nM, 1 h). Then, nuclei were stained using Hoechst 33342 (62249;Invitrogen, 10mg/mL) at 1:2000 dilution for 30 mins. Cells were rinsed twice with 1xPBS and imaged in phenol-free DMEM with DMSO or apilimod. Images were acquired with a Nikon Ti2-E inverted microscope equipped with a Yokogawa CSU-W1 SoRa spinning-disk super-resolution module using a 60× SR Plan Apo IR oil-immersion objective. Images were processed and adjusted in FIJI (version 2.14.0/1.54f). Average lysosome volume per cell was measured using Imaris software (version 10.2.0). A detailed protocol is available at https://dx.doi.org/10.17504/protocols.io.6qpvr1d5bgmk/v1.

### siRNA Transfection

siRNA-mediated knockdowns of target gene expression were accomplished using Horizon Biosciences siGENOME-pooled siRNAs. Briefly, 2.5 × 105 RAW 264.7 cells were plated per well in a 6-well dish. The following day, cells were transfected using Lipofectamine RNAiMAX (2448190; Invitrogen) as per the manufacturer’s protocols using a 100 nM siRNA pool. After 48 h, cells were treated with drugs or lysed and subjected to immunoblotting. A detailed protocol can be found at: https://dx.doi.org/10.17504/protocols.io.4r3l29owjv1y/v1.

### Statistical analysis

All quantitative data were obtained from at least three independent biological replicates unless otherwise indicated. Immunoblot signals were quantified using FIJI (ImageJ, version 2.14.0/1.54f). Data are presented as mean ± SEM. Statistical analyses were carried out using GraphPad Prism (10.3.1). Comparisons between two groups were performed using two-tailed unpaired Student’s *t* tests. Multiple group comparisons were analyzed by one-way ANOVA with Sidak’s post-test. The number of replicates (*n*), type of statistical test, and *P* values for each experiment are reported in the corresponding figure legends.

## Acknowledgements

We are grateful to all members of the Ferguson laboratory and to Pietro De Camilli. This research was supported by grants from the NIH (GM105718), the Carol & Gene Ludwig Program for the Study of Neuroimmune Interactions in Dementia at Yale University (Ludwig Development Award to S.F.), Aligning Science Across Parkinson’s disease (ASAP-000580) through the Michael J. Fox Foundation for Parkinson’s Research (MJFF) to S.F.. We thank Berrak Ugur (Yale) and Benjamin Johnson (Yale) for managing our ASAP project. We thank Fiona Weaver (Yale) for help with designing the summary diagram in Figure 5. The authors declare no competing financial interests.

## Author Contributions

D.C., A.B.D. and S.F. conceptualized the project. D.C. performed most of the experiments. A.B.D. investigated link between CASM and LRRK2 activation and made key observations that were critical for initiation of the project. A.R.F. performed experiments on human microglia. All authors edited and provided comments on the manuscript.

## Data Availability

All data, protocols, and key laboratory materials used or generated in this study are listed in the Key Resources Table, together with their persistent identifiers (10.5281/zenodo.22117758). No new code was generated for this study.

