## Supplemental Data for "ClC-7 links PIKfyve inhibition to Rab-dependent LRRK2 activity at lysosomes"

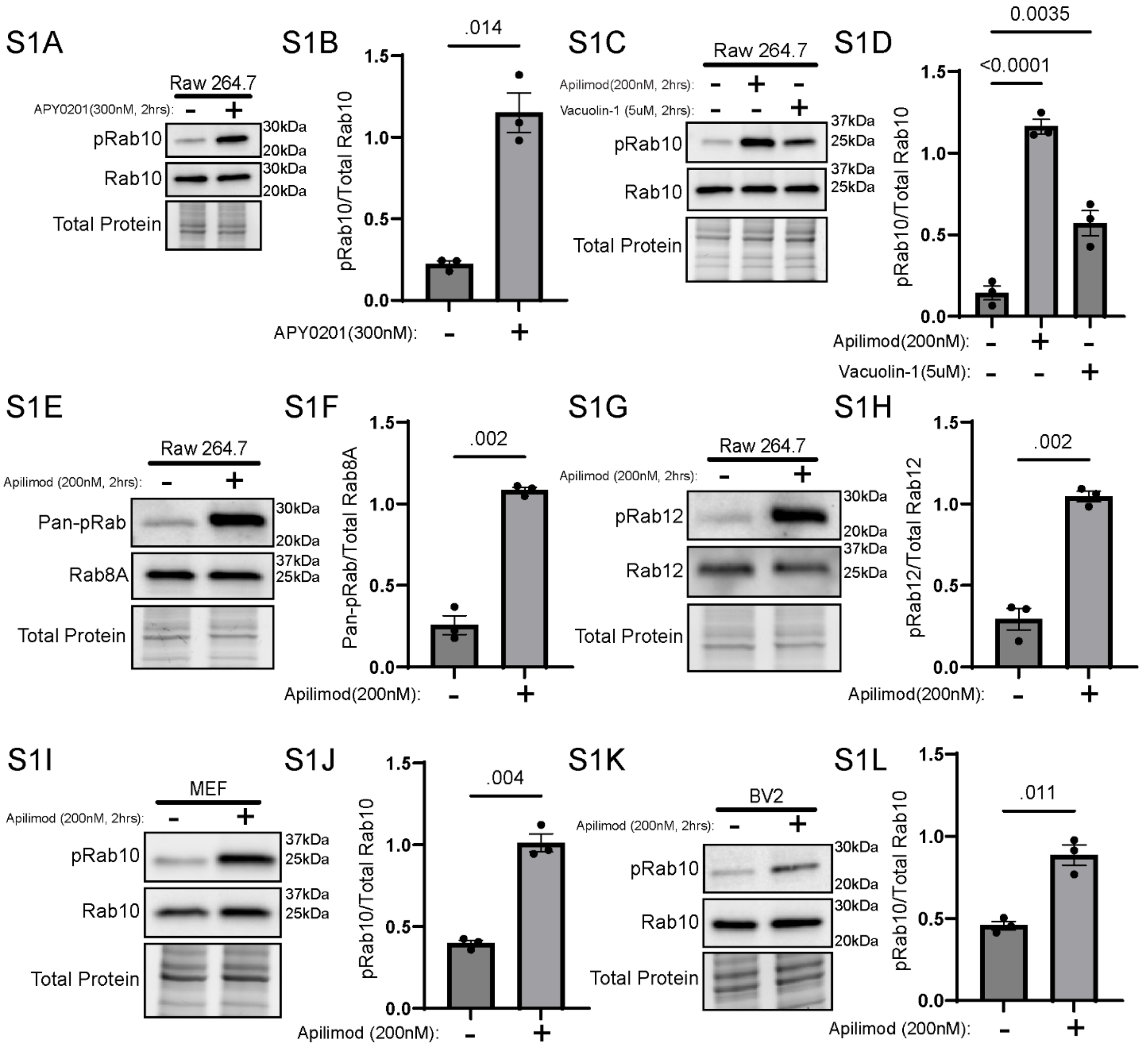


**Supplemental Figure 1. Validation of PIKfyve-dependent LRRK2 activation across inhibitors and cell types.** (A) Immunoblot of Raw 264.7 cells treated with APY0201 [300nM for 2 h, pRab10 (T73)]. (B) Quantification of pRab10 (T73) after APY0201 (mean ± SEM; raw values, n = 3, two-tailed unpaired Welch's t test). (C) Immunoblot of Raw 264.7 cells treated with apilimod (200nM, 2 h) or vacuolin-1 (5µM, 2 h) and probed for pRab10 (T73). (D) Quantification of pRab10 (T73) after apilimod or vacuolin-1 treatment (mean ± SEM; raw values, n = 3, one-way ANOVA, Sidak's). (E–H) Immunoblots and quantifications of pRab8 (T72) and pRab12 (S106) after apilimod treatment (mean ± SEM; raw values, n = 3, two-tailed unpaired Welch's t test. (I–L) Immunoblots and quantifications of MEFs and BV-2 microglia showing elevated pRab10 (T73) after apilimod (mean ± SEM; raw values, n = 3, two-tailed unpaired Welch's t test).

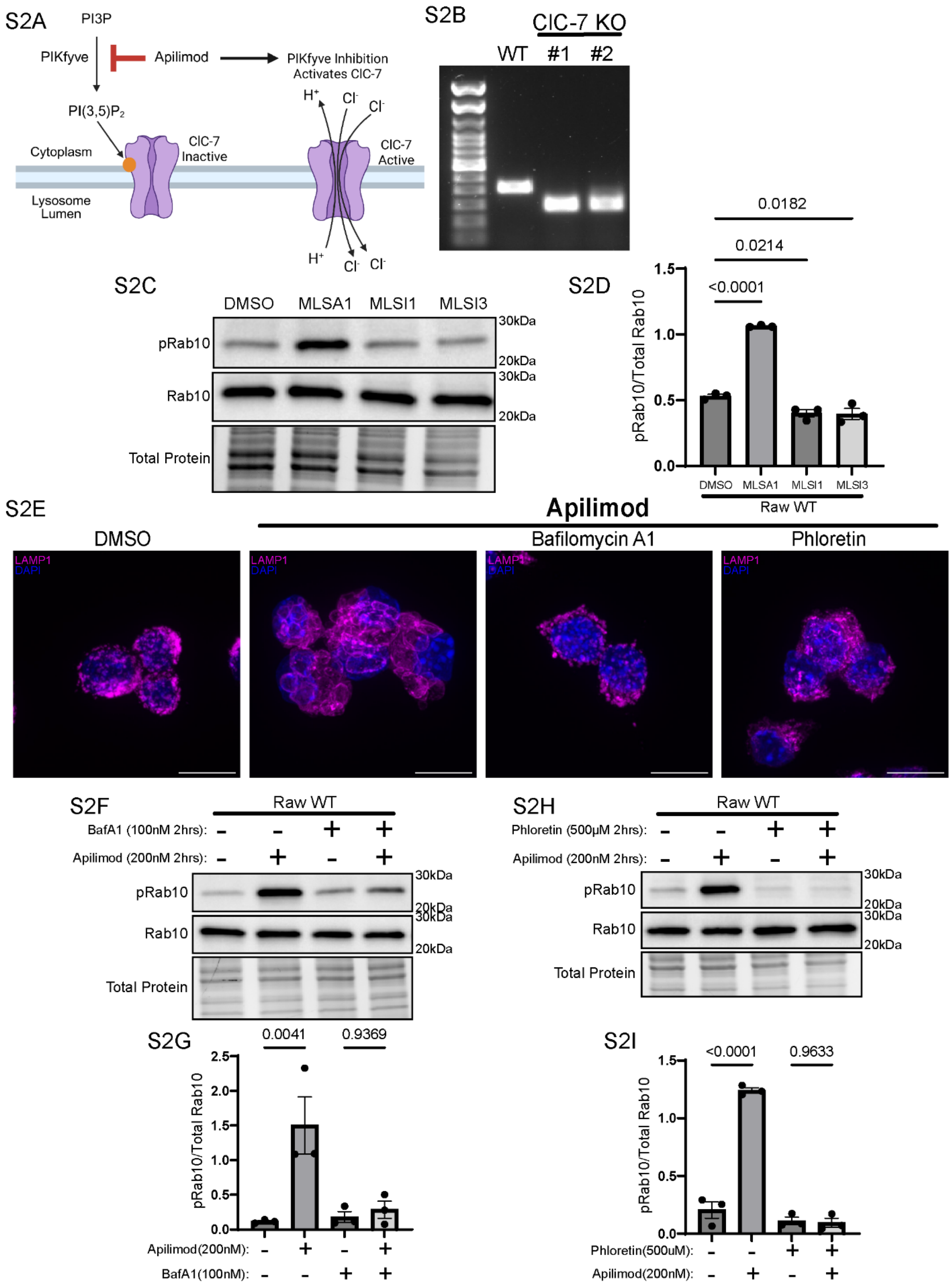
S2A
PI3P
PIKfyve
Apilimod
PIKfyve Inhibition Activates ClC-7
PI(3,5)P2
Cytoplasm
ClC-7 Inactive
Lysosome Lumen
ClC-7 Active
S2B
ClC-7 KO
WT
#1
#2
S2C
DMSO
MLSA1
MLSI1
MLSI3
pRab10
Rab10
Total Protein
30kDa
20kDa
S2D
pRab10/Total Rab10
<0.0001
0.0214
0.0182
Raw WT
S2E
Apilimod
DMSO
Bafilomycin A1
Phloretin
LAMP1
DAPI
S2F
Raw WT
BafA1 (100nM 2hrs):
Apilimod (200nM 2hrs):
S2G
0.0041
0.9369
Apilimod(200nM):
BafA1(100nM):
S2H
Phloretin (500μM 2hrs):
S2I
<0.0001
0.9633
Phloretin(500uM):
Apilimod(200nM):

**Supplemental Figure 2. Clcn7 knockout validation, TRPML1 modulation of LRRK2 activity, and Bafilomycin and Phloretin Block Enlarged Lysosomes and LRRK2 Activation after PIKfyve Inhibition.** (A) Schematic showing PIKfyve regulates ClC-7 ion transporter activity through interaction with PI(3,5)P2. Inhibiting PIKfyve using apilimod causes a loss of PI(3,5)P2 and an increase in ClC-7 activity. Created with BioRender (Clegg, D. and Ferguson, S.M., 2026). (B) 1 % agarose gel confirming Clcn7 KO Raw264.7 clones. (C) Immunoblot analysis of cells treated with TRPML1 agonist ML-SA1 or inhibitors ML-SI1 and ML-SI3 (10uM, 20uM, and 25uM, respectively) and probed for pRab10 (T73). (D) Quantification of pRab10 (T73) after ML-SA1, ML-SI1, or ML-SI3 (mean ± SEM; raw values, n = 3, one-way ANOVA, Sidak's). (E) Immunofluorescent images of Raw264.7 cells DMSO, Apilimod (200nM, 2 h), Apilimod + Bafilomycin A1 (100nM, 2 h), or Apilimod + Phloretin (500μM, 2 h) (LAMP1, magenta; DAPI). (F) Immunoblot analysis of cells treated with Apilimod (200nM, 2 h) ± Bafilomycin A1 (100nM, 2 h) and probed for pRab10 (T73). (G) Quantification of pRab10 (T73) in cells treated with Apilimod ± Bafilomycin A1 (mean ± SEM; raw values, n = 3, one-way ANOVA, Sidak's). (H) Immunoblot analysis of cells treated with Apilimod (200nM, 2 h) ± Phloretin (500μM, 2 h) and probed for pRab10 (T73). (I) Quantification of pRab10 (T73) in cells treated with apilimod ± Phloretin (mean ± SEM; raw values, n = 3, one-way ANOVA, Sidak's).

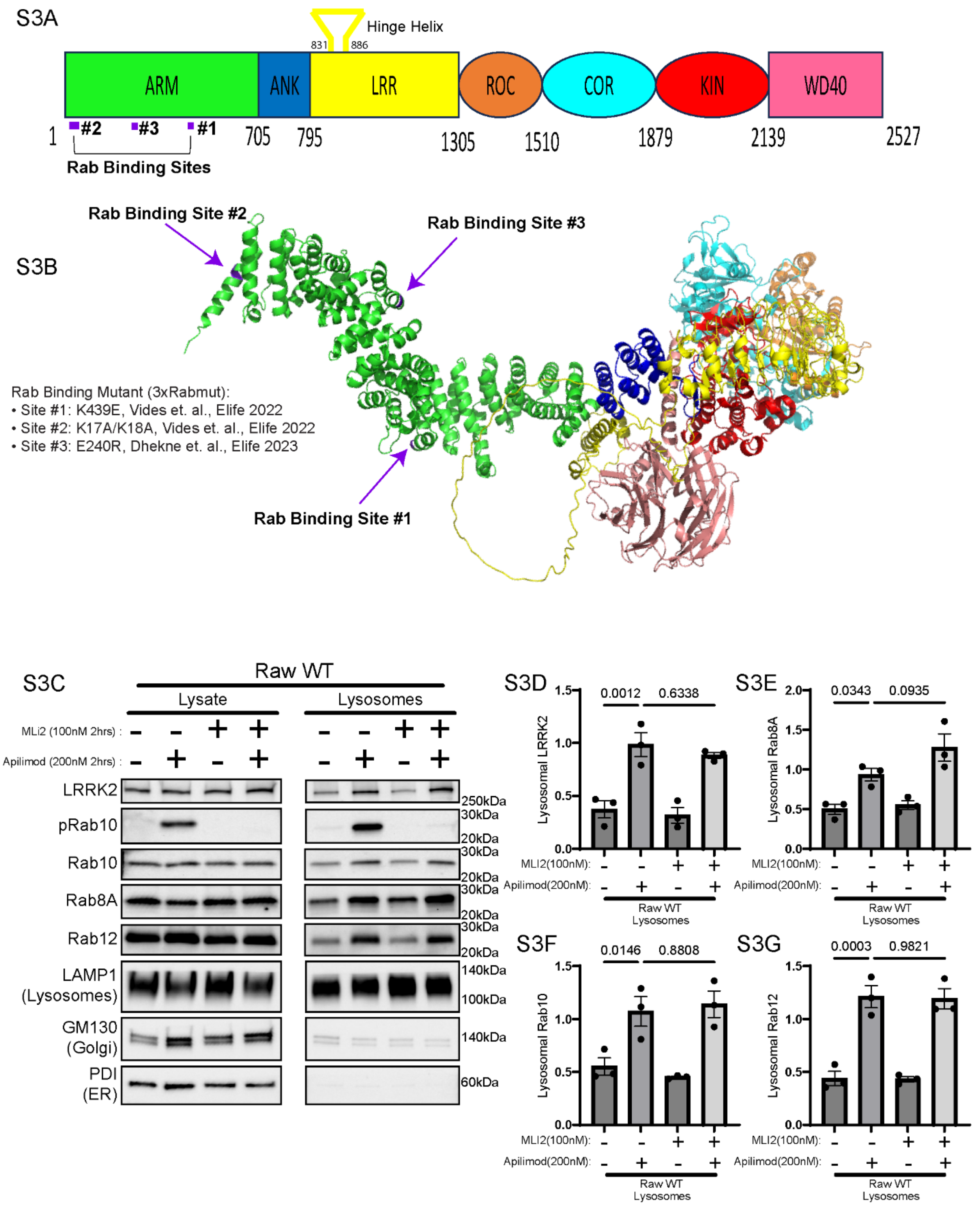
S3A
Hinge Helix
831
886
ARM
ANK
LRR
ROC
COR
KIN
WD40
1
#2
#3
#1
705
795
1305
1510
1879
2139
2527
Rab Binding Sites
S3B
Rab Binding Site #2
Rab Binding Site #3
Rab Binding Site #1
Rab Binding Mutant (3xRabmut):
• Site #1: K439E, Vides et. al., Elife 2022
• Site #2: K17A/K18A, Vides et. al., Elife 2022
• Site #3: E240R, Dhekne et. al., Elife 2023
S3C
Raw WT
Lysate
Lysosomes
MLi2 (100nM 2hrs) :
Apilimod (200nM 2hrs) :
LRRK2
pRab10
Rab10
Rab8A
Rab12
LAMP1 (Lysosomes)
GM130 (Golgi)
PDI (ER)
250kDa
30kDa
20kDa
30kDa
20kDa
30kDa
20kDa
30kDa
20kDa
140kDa
100kDa
140kDa
60kDa
S3D
0.0012
0.6338
Lysosomal LRRK2
S3E
0.0343
0.0935
Lysosomal Rab8A
S3F
0.0146
0.8808
Lysosomal Rab10
S3G
0.0003
0.9821
Lysosomal Rab12
MLI2(100nM):
Apilimod(200nM):
Raw WT Lysosomes

**Supplemental Figure 3. Rab binding is required for LRRK2 recruitment but not for Rab accumulation.** (A) Sequence of LRRK2 showing its three Rab-binding sites. (B) AlphaFold-predicted LRRK2 structure showing three Rab-binding regions (purple arrows). (C) Immunoblot lysates and lysosomes isolated from wild-type cells treated with apilimod (200nM, 2 h) ± MLi-2 (100nM, 2 h). Images for the lysate and lysosome fractions come from the same exposure of the same blot that was loaded with 5μg lysate and 0.5μg of lysosome protein. (D–G) Quantification of lysosomal LRRK2 and Rabs (Rab8, Rab10, Rab12) after MLi-2 ± apilimod (mean ± SEM; raw values, n = 3, one-way ANOVA, Sidak's).
